# Left-right alternation of superior colliculus activity with the stepping rhythm

**DOI:** 10.64898/2026.08.18.745617

**Authors:** Cameron Wilhite, Loren M. Frank, Massimo Scanziani

## Abstract

Fluid behavior requires temporal coordination across brain systems that orchestrate movement. A clear example is locomotion, in which left and right turns are not only precisely coordinated with the ongoing stepping rhythm but occur at opposite phases of the stepping cycle. The circuits underlying this coordination remain poorly understood. We discover that neuronal activity in the mouse superior colliculus, a conserved midbrain structure involved in turning behavior, is tightly phase-locked to the stepping rhythm. Notably, neurons selective for left and right turns fire at opposite phases of the stepping cycle. Moreover, this phase opposition is already evident during straight locomotion, before the animal initiates a turn. By aligning the activity of left and right turn neurons to opposite phases of the stepping cycle, the superior colliculus may create alternating windows of opportunity for left and right turns, facilitating the seamless execution of turns during locomotion.

## Introduction

The precise temporal coordination of distinct movements relative to each other is essential for fluid behavior. Even simple behavior like turning left or right during locomotion relies on the precise orchestration of movements. While such turns unfold seamlessly along the animal’s trajectory, they are in fact tightly coordinated with the ongoing locomotor stepping rhythm^1,2^. If humans walking along a straight path are cued to turn in a direction of their choosing, for example, the direction will depend on which foot is planted around the time of the cue^1,2^. Thus, opposite phases of the stepping cycle favor turns in opposite directions. Coordinating turns with specific phases of the stepping cycle ensures that the animal’s center of mass is optimally supported throughout the turn while also minimizing the torque required to rotate the body^1–3^ . The stability and efficiency of turns obtained through this coordination are likely fundamental during pursuit and escape behaviors. The identity of the circuits that maintain this coordination is, however, not well understood.

We addressed this question by recording neuronal activity from the superior colliculus (SC), a conserved vertebrate midbrain structure that plays a central role in turning behavior^4,5^, in freely moving mice. The intermediate and deep motor layers of the SC (dSC) contain “turn cells” whose activity reports turn direction^6–10^ and whose stimulation triggers turning movements^11–13^. To determine whether turn-related activity in these neurons is coupled to the stepping rhythm we recorded from the dSC while monitoring locomotor dynamics of mice engaging in left or right turns. We discover that the activity of a large population of turn cells is profoundly modulated by the stepping rhythm and that their turn direction preference, whether left or right, dictates the stepping phase during which they fire most. Consequently, left and right turn cells fire preferentially at opposite phases of the stepping rhythm, an alternation which is apparent already during straight locomotion. These results reveal how turn related activity in the superior colliculus, one of the best understood motor functions of this structure, unfolds in tight temporal coordination with the ongoing behavior of the organism.

## Results

### Coordination of turn direction with step cycle

The coordination between turn direction and the step phase is well established in humans^1,2^, but whether this also occurs in mice has not been verified. We therefore assessed the stepping phase of mice as they performed left or right turns. We first characterized the stepping rhythm of mice on a transparent linear track using a bottom-view camera. As in rodents^14^ and other walking tetrapods^15,16^, with each forepaw swing, the head shifted laterally such that a right forepaw swing was linked to a leftward head movement and vice versa, producing a left-right oscillation of the head during locomotion (Figure 1A and Figure S1). This allowed us to infer the stepping rhythm of mice by monitoring the left-right oscillation of the head with a top-view camera, quantifying this oscillation as the cyclic change in head-body angle (Figure 1B, left, See Methods). We subdivided this oscillation into the following phases of the stepping cycle (Figure 1B, right panel): phase π/2, when the head, aligned straight with the body, is moving left and the right forepaw is swinging forward, phase 3π/2 when the head, aligned straight with the body, is moving right and the left forepaw is swinging forward and phases 0 and π, when the animal’s head is pointing to the right or left, respectively, relative to the body. To obtain consecutive trials in which the animals alternated between left and right turns, we trained mice on a Y-maze spatial navigation task (Figure 1C) in which two of the three reward ports (located at the ends of the maze arms) were active, with their locations changing uncued across blocks. During each trial, the mouse shuttled from one reward port to another, resulting in repeated left and right turns at the maze bifurcation. During locomotion on the Y-maze, the amplitude of the head-body angle oscillated between positive and negative ∼2.5 degrees (2.64 ± 0.06 deg., average ± s.e.m.; here and throughout unless stated otherwise) at ∼5.5 Hz (5.45 ± 0.02 Hz, ∼11 steps per second). Crucially, animals reached the center point of the maze bifurcation at opposite phases of the stepping cycle depending on the direction of the turn (Figure 1D, left turns near phase 0 (2π), 6.2 ± 0.1 rad., right turns near phase π, 2.9 ± 0.1 rad., n = 55 sessions, p < 0.001 for each turn direction, Rayleigh tests, p < 0.05, paired Rayleigh test, mean phase difference: 3.0 rad.). Thus, at the maze bifurcation, left and right turns occur at opposite phases of the stepping cycle, consistent with other organisms^1,2^.

**Figure 1:**
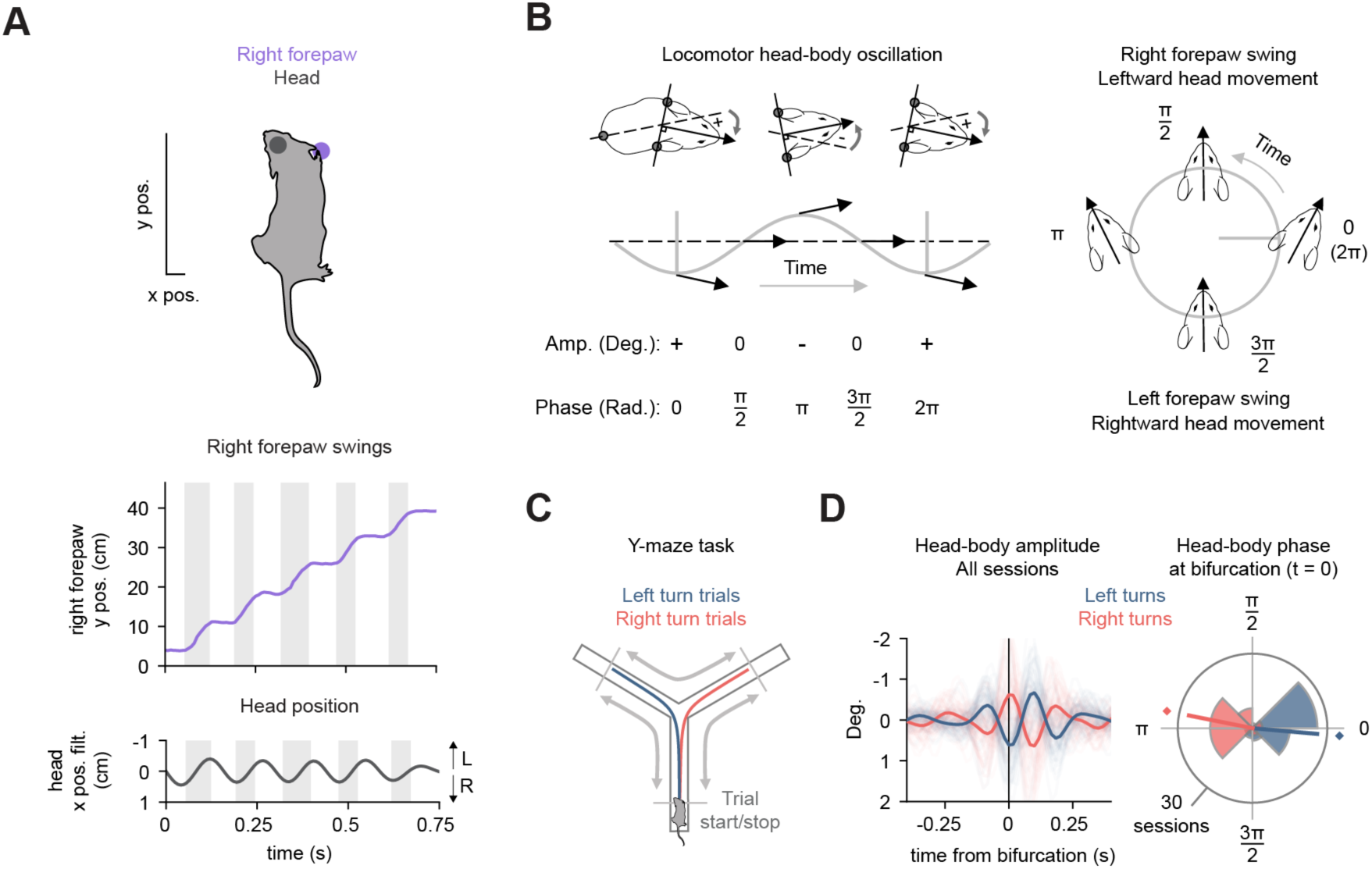
Turns are coordinated with the stepping cycle in mice. **(A)** Above, schematic of mouse with labels on the right forepaw and head (as labeled on frames taken with the bottom view camera; scale bar: 1 cm x, 8 cm y). Below, example pass on linear track. Top panel: Right forepaw position along the y-axis (y pos.). Gray bands indicate times of right forepaw swings. Bottom panel: Head position along the x-axis (x pos.) filtered between 4-10 Hz. Note coupling between right forepaw swings and leftward head movements. **(B)** Left, schematic illustration of head-body angle during locomotion. Positive and negative amplitude values reflect right and left head direction relative to the body, respectively. The phase of the head-body oscillation cycle is given in radians. Right, polar plot showing head and forepaw movements associated with each quarter phase of the head-body oscillation cycle. **(C)** Y-maze with average behavioral trajectories of left and right turns across all sessions and mice overlaid (n = 55 sessions, 12 mice). The crossing of the 70% point on the maze arms (gray lines) determines the start and stop of each trial. **(D)** Left, amplitude of head-body angle filtered between 4-8 Hz around the time animals reached the bifurcation across all sessions color coded by turn direction. Time from bifurcation was defined as the time at which the animal’s head position crossed the center point of the maze. Dark lines: average over all sessions and animals (n = 55 sessions, 12 mice); light lines: individual sessions. Note similar amplitude yet opposite sign of the head-body angle around the bifurcation. Right, distributions of instantaneous head-body phase at the time the animals reached the bifurcation (t = 0) for each turn direction across sessions (left turns: p < 0.001 Rayleigh test, right turns: p < 0.001, Rayleigh test. Paired Rayleigh test between left and right turns: p < 0.05).

### Turn cells in the dSC are locked to the step cycle

Is the activity of turn cells in the dSC modulated in coordination with the stepping cycle and, if so, are left and right turn cells active at opposite phases of the cycle? To address this question, we used silicon probes to record from dSC neurons (Figure 2A, top panel, Figure S2) as mice performed the Y-maze task (Figure 2A, middle panel). We identified turn cells as neurons whose activity was differentially modulated for left or right turns at the maze bifurcation using the Turn Direction Selectivity Index (TDSI; See Methods), where negative or positive values correspond to left or right turn cells, respectively. Turn cells were defined as cells whose TDSI maintained the same sign and remained significant regardless of which maze arm the animal approached the bifurcation from (Figure 2B and 2C, 28% of cells, 411/1471 from 12 animals, TDSI for each arm: p < 0.05, permutation tests). Turn cell firing peaked around the bifurcation (+18 ± 7 ms, Figure 2D, bottom panel) and became directionally selective ∼300 ms (283 ± 11 ms) before the bifurcation (Figure 2D, top panel) with a subset (51%, 211/411) achieving selectivity prior to the time at which the animals’ behavior diverged between left and right turns (Figure 2A, bottom panel). Consistent with the lateralization of the dSC hemispheres for left and right turns^8,9,17,18^, the majority of turn cells showed selective firing for contraversive turns (Figure 2E, contraversive (e.g. left turn cells in the right dSC): 69%, 283/411, ipsiversive (e.g. right turn cells in the right dSC): 31%, 128/411). Together, the properties of the turn cells identified here are consistent with the reported characteristics of motor command neurons responsible for turns in the dSC^8–11^.

**Figure 2:**
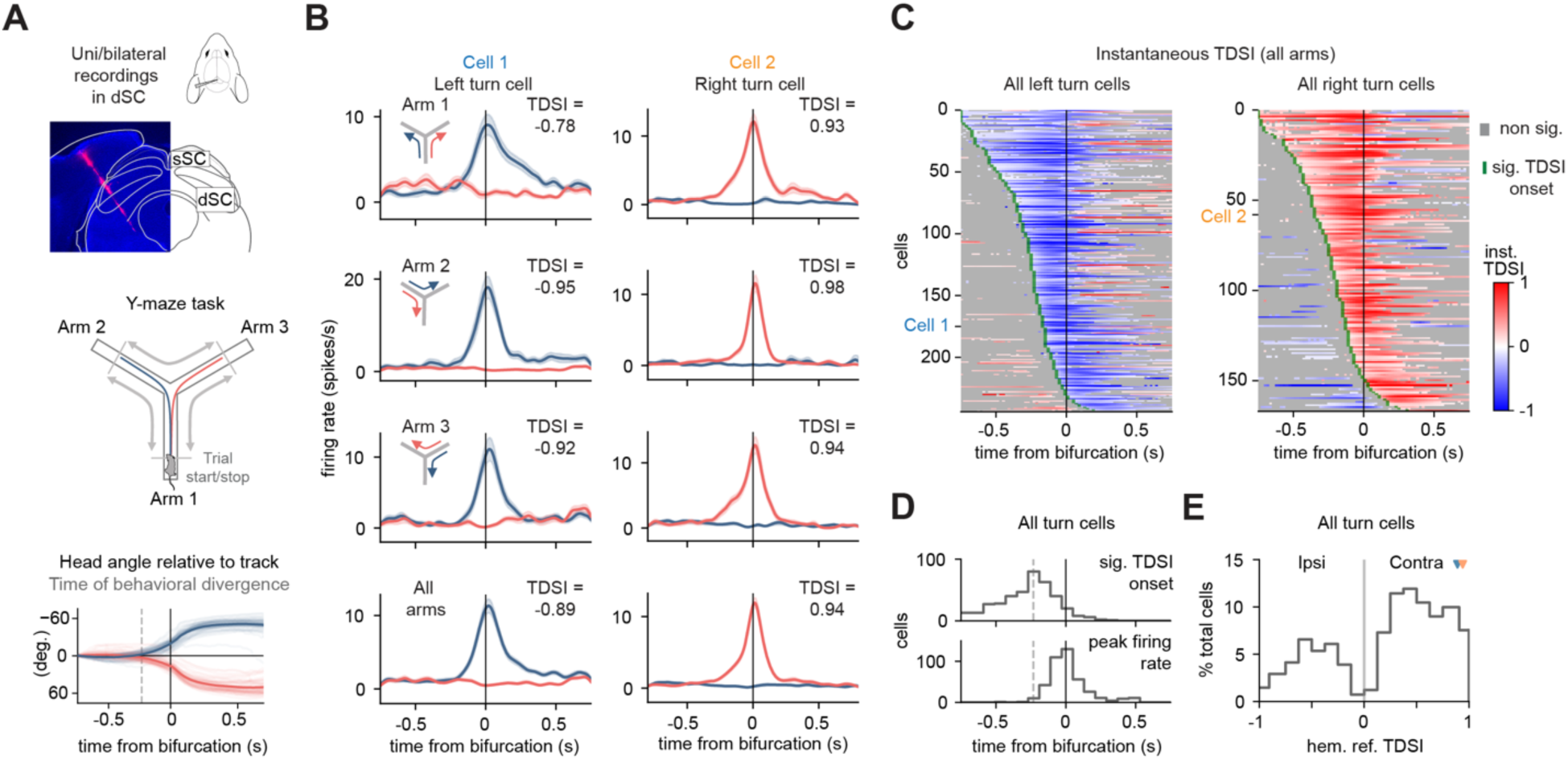
Turn cells in the intermediate and deep layers of the SC. **(A)** Recording configuration and Y-maze task. Top, schematic and histology of an example implantation targeting the intermediate and deep layers of the SC (dSC). Probe track is marked by DiI (pink). sSC = superficial layers of the SC. Middle, Y-maze with behavioral trajectories. Bottom, angle of the animals’ head relative to the track during the task. Dark lines: average over all sessions and animals (n = 55 sessions, 12 mice); light lines: individual sessions. Dashed line: time from the maze bifurcation at which the average head angle relative to the track significantly diverges between right and left turns (-0.25 s). **(B)** Firing rates of two example turn cells for all possible trajectories on the Y-maze. Left, example left turn cell (Cell 1). Right, example right turn cell (Cell 2). Blue and red are left and right turn trajectories, respectively. Bottom panels are the averages of the firing rates across all left or right turn trajectories. Thick lines: average firing rates across trials, shaded bands represent s.e.m. Note negative and positive TDSI (Turn Direction Selectivity Index) values for Cell 1 and 2 respectively, across all arms (TDSI: p < 0.001 for each arm, permutation tests). **(C)** All left and right turn cells sorted by the time at which the instantaneous TDSI became selective in the cell’s preferred turn direction (significant TDSI onset; n = 244 left turn cells, n = 167 right turn cells; 8 right dSC insertions, 34 sessions; 6 left dSC insertions, 28 sessions). **(D)** Distributions of significant TDSI onset times and times of peak firing rate across all turn cells. Dashed lines: average time of behavioral divergence as shown in panel A. **(E)** Distribution of TDSI values across all turn cells referenced to the recorded dSC hemisphere. Positive and negative hemisphere referenced TDSI values indicate turn cells whose firing is selective for contraversive (e.g. left turn cells in the right dSC) or ipsiversive (e.g. right turn cell in right dSC) turns, respectively. Markers indicate hemisphere referenced TDSI values for Cell 1 and Cell 2.

We discovered that nearly half of the turn cells (47%, 195/411) exhibited firing that was significantly phase-locked to the stepping cycle (Figure 3A, example cells). This phase-locked firing was evident throughout the maze, both during straight locomotion as well as during the turn, that is, also outside of periods when turn cells showed direction-selective firing. Strikingly, left and right turn cells fired at opposite phases of the stepping cycle (Figure 3B). For the most accurate assignment of the preferred phase at which turn cells fired, we focused on those cells with an absolute TDSI of at least 0.1 and whose activity was most strongly modulated in coordination with the stepping cycle (i.e., with a ln(Rayleigh Z) > mean + SD of population, n = 49 cells, Figure S3, See Methods). We found that left turn cells fired preferentially near phase π/2, that is, when the right forepaw is swinging forward and the head is moving left relative to the body (1.70 ± 0.23 rad., n = 31 cells). In contrast, right turn cells fired near the opposite phase, 3π/2, that is, when the left forepaw is swinging forward and the head is moving right relative to the body (4.21 ± 0.28 rad., n = 18 cells; Figure 3C, p < 0.05 for both left and right turn cell populations, Rayleigh tests for non-uniformity). Preferred firing phases were the same regardless of the direction of the upcoming turn (Figure S4A, *rc* = 0.902, p < 0.001, permutation test).

**Figure 3:**
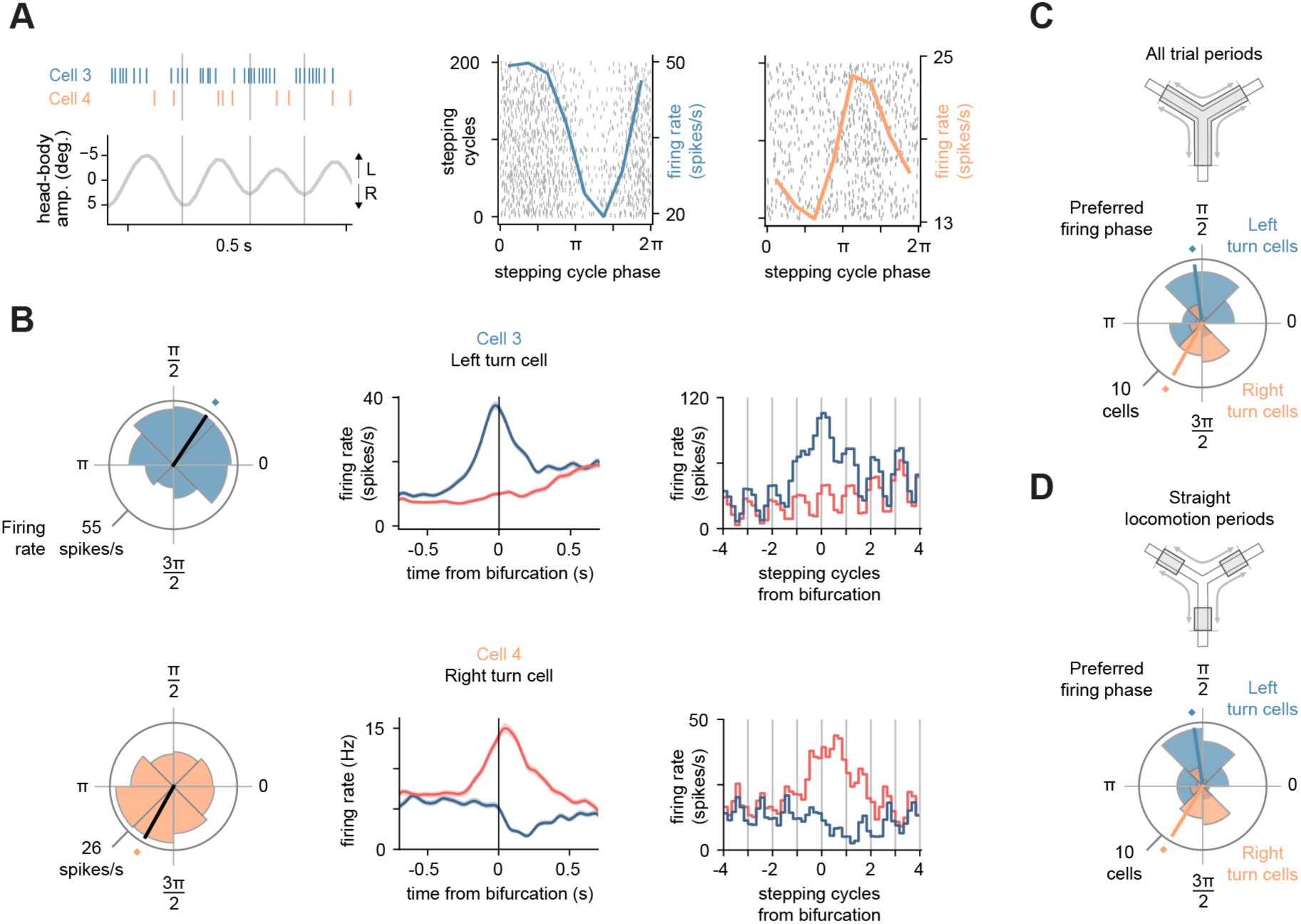
Turn cell activity is coupled to the stepping rhythm. **(A)** Left, spike raster of two simultaneously recorded turn cells, Cell 3 and Cell 4 (top) and ongoing head-body oscillation (bottom). Vertical gray lines indicate phase 0 (2π). Middle and right, spike raster plots of Cell 3 and Cell 4, respectively, aligned to the first 200 head-body oscillation cycles in the session. The overlaid traces are the firing rates averaged over all head-body oscillation cycles in the session (right y-axis). Note that the firing of these two cells is coupled to opposite phases of the head- body oscillation cycle. **(B)** Left, polar plots of the firing of Cell 3 and Cell 4 relative to the phase of the head-body oscillation cycle for the session shown in panel A. Cell 3: Ln(Z): 7.15, p < 0.001, Rayleigh test, preferred firing phase: 0.98 radians; Cell 4: ln(Z): 5.30, p < 0.001 Rayleigh test, preferred firing phase: 4.20 radians. Middle, firing rates of Cell 3 and Cell 4 around the time of left (blue) and right (red) turns showing their opposite selectivity for turn direction (Cell 3: TDSI = -0.55; Cell 4: TDSI = 0.56). Right, firing rates of Cell 3 and Cell 4 aligned to head-body cycles from turn. Vertical gray lines indicate phase 0 (2π). **(C)** Preferred firing phases of left and right turn cells during all trial periods (cells with an absolute TDSI of at least 0.1 and ln(Rayleigh Z) > mean + SD of population; n = 49 total cells, 31 left turn cells, 18 right turn cells). Left and right turn cell populations were significantly clustered around phases 1.70 and 4.21 rad., respectively (left turn cell population: p < 0.01, right turn cell population: p < 0.05, Rayleigh tests) and significantly different from one another (left vs. right turn cell populations: p < 0.05, permutation test, mean phase difference: 2.51 rad.). **(D)** Same as in C but during straight locomotion periods only. Left and right turn cell populations were significantly clustered around phases 1.72 and 4.16 rad., respectively (left turn cell population: p < 0.01, right turn cell population: p < 0.05, Rayleigh tests) and significantly different from each other (left vs. right turn cell populations: p < 0.05, permutation test, mean phase difference: 2.44 rad.).

Importantly, these cells maintained the same phase preference also during periods of straight locomotion, before and after the animal engaged in the turn (Figure S4B, *rc* = 0.998, p < 0.001, permutation test). The preferred firing phases of left and right turn cells were significantly different both across the population (Figure 3C, all trial periods: p < 0.05, permutation test, mean phase difference: 2.51 rad.; Figure 3D, straight locomotion periods: p < 0.05, permutation test, mean phase difference: 2.44 rad.; Figure S4C, distribution across animals) and in the phase differences calculated across simultaneously recorded cell pairs (Figure S4D). Notably, the extent of the phase separation between left and right turn cells varied as a function of the strength of their modulation with the stepping cycle (i.e. the Rayleigh Z). Lowering the Rayleigh Z threshold to include more weakly phase-locked cells progressively reduced the phase separation, whereas raising the threshold increased it (Figure S5), indicating that the phase separation between left and right turn cells is most pronounced among turn cells with the strongest rhythmic activity.

In summary, during locomotion, SC turn cell activity is rhythmically coupled to the ongoing stepping rhythm, such that left and right turn cells fire at opposite phases of the stepping cycle. Accordingly, the turn direction selectivity of a rhythmically active SC cell is revealed not only around the turn but also during straight locomotion, reflected in its preferred firing phase relative to the stepping cycle.

## Discussion

Our study shows that turn-related activity in the dSC, possibly among the best understood motor outputs of this structure, is tightly coupled to ongoing locomotor behavior, such that left and right turn cells fire preferentially at opposite phases of the stepping cycle. To our knowledge this is the first report of oscillatory neuronal activity in the dSC that is coupled to the stepping rhythm. Because of the opposite phases of the stepping cycle at which left and right turn cells fired, their direction preference could be inferred also during straight locomotion. The alignment of left and right turn cell activity to opposite phases of the stepping cycle is likely to create windows of opportunity that alternatively favor left or right turns. This alignment may underlie the precise coordination between turn direction and stepping phase observed in humans^1,2^ and validated here in mice (Figure 1D).

The turn cells identified in this study share several characteristics with previously described motor command neurons for orienting movements in the SC: 1) turn cells are located in SC’s intermediate and deep layers, layers whose neurons project to brainstem turning centers^19,20^, and whose stimulation elicits turning movements^11–13,^ 2) turn cell activity primarily encodes contraversive turning movements, consistent with dSC motor signals^8,9,17,18^, 3) turn cells become directionally selective before the time at which the animal’s behavior diverges between left and right turns, consistent with premotor activity in the dSC^8,9,17,18,21^, and 4) turn cells fire similarly at the maze bifurcation regardless of the animal’s starting position, indicating that turn cell activity reflects an egocentric, displacement-based motor response. These observations suggest that the turn cell activity reported here primarily reflects a motor command for turning.

Locomotor rhythmic activity in dSC turn cells could therefore reflect active motor commands that produce small, left-right head movements during each stepping cycle. Indeed, left turn cells fired preferentially near the phase of the stepping cycle when the head is moving left relative to the body, and right turn cells fired near the phase when the head is moving right (Figure 3C). Executing a larger turn during locomotion could be achieved by “superimposing” extra excitation onto motor command neurons that are already phasically active on each step^22^. Alternatively, locomotor rhythmic activity in turn cells could reflect sensory^23–26^ or motor-related^27–29^ inputs that oscillate with stepping. Because not all turn cells exhibited activity that was phase-locked to the stepping cycle, the neurons described in this study may constitute a distinct cell type with specific connectivity patterns.

In our companion paper^30^, we show that dSC turn cells can also be modulated in coordination with hippocampal representations of possible future paths (i.e., hippocampal sweeps). Because approximately half of the dSC turn cells whose activity was modulated in coordination with hippocampal sweeps also exhibited activity that was significantly phase-locked to the stepping cycle (48%, 27/56 cells), these two forms of modulation of turn cell activity are not mutually exclusive.

In conclusion, this study reveals the phase locking of turn-related activity in the dSC to the stepping rhythm. This temporal modulation ensures the coordination between motor commands for a specific movement with the ongoing behavior of the organism as it navigates through its world.

## Methods

### Experimental model and animals

Neural activity was recorded from twelve male C57BL/6J mice (Jackson Laboratory #000664, 3.5- 8 months old). All mice were housed on a reversed cycle (light/dark cycle 12/12h). Mice were housed with littermates before experimentation and singly housed in enriched cages during food-restriction and experimental protocols. All experimental procedures were conducted in accordance with the regulations of the University of California San Francisco Institutional Animal Care and Use Committee (IACUC).

### Behavioral training and electrode implantation

The implantation procedure was performed in two phases on two distinct days. First, the implantation of the headplate, and second, the implantation of the silicon probe(s). Mice were implanted under 2% isoflurane anesthesia. The headplate was placed on the skull with dental cement (Unifast LC, GC America; Optibond Universal, Kerr Dental). After post-operative recovery of five days in their home cage, mice were deprived of food to 85% of their baseline weight and pretrained to run on an elevated 1 m linear track for liquid reward (sweetened evaporated soymilk). This training was done to familiarize the mice with running on a track and accessing reward ports. After the mice alternated between the reward ports reliably on the linear track (1-3 days), they were pretrained on the Y-maze task (3-5 days, see *Y-maze task and trial segmentation*). After successful pretraining on the Y-maze, mice underwent the silicon probe implantation surgery. A silicon probe was mounted on a 3D-printed moveable microdrive to record unilaterally or bilaterally from the superior colliculus (SC). In five of the twelve mice, a silicon probe was also implanted in the dorsal CA1 region of the hippocampus to collect the data presented in our companion paper ^30^. Only SC data were analyzed in this study. The types of probes used included: Single-shank probes (Diagnostic Biochips (DBC) P64-4, 64 channels per shank, 20 μm interchannel pitch), four-shank probes (DBC P64-1, 250 μm inter-shank distance, 16 channels per shank, 20 μm interchannel pitch, DBC P128-6, 150 μm inter-shank distance, 32 channels per shank, 25 μm interchannel pitch), and eight-shank probes (DBC P128-8, 150 μm inter-shank distance, 16 channels per shank, 30 μm interchannel pitch). Probe shanks were inserted through a small craniotomy at the coordinates described in the *Electrode insertion, electrode coordinates, histology and recording site assessment* section below. The shanks were initially lowered to 0.7-1.2 mm below the surface of the brain and the microdrive was then fixed, via dental cement, to the headplate. A ground electrode (0.005” diameter stainless wire, A-M systems) was inserted in the cerebellum. After post-operative recovery, mice were placed back on the Y-maze training protocol for 1-5 days while the shanks were lowered to the intermediate and deep SC (dSC). This protocol ensured that peak Y-maze performance coincided with the time at which the recording sites reached the dSC. Mice remained on food restriction for the remainder of the experiment to ensure engagement with the Y-maze task.

### Electrophysiological recordings and video tracking

The data presented in this paper are from three to eight 45-120 min sessions per mouse on the Y-maze task. For the stepping analysis, two implanted mice also ran on a transparent linear track (Figure 1, Figure S1, see *Transparent linear track task and stepping analysis*). Electrophysiological data were acquired using an Intan RHD2000 system (Intan Technologies LLC) band-pass filtered between 0.1 Hz and 7.5 kHz and digitized at 20 kHz. Spike sorting was performed semi-automatically, using Kilosort 2.0^31^. This was followed by manual adjustment of the waveform clusters using the software Phy. For the Y-maze task, video data was acquired at 60 frames per second using a CMOS camera (Basler acA1300-200um) placed above the maze. For the transparent linear track task, video was acquired at 124 frames per second using a CMOS camera (Basler acA1300-200um) with a wide-angle lens placed below the track. A machine-learning algorithm, DeepLabCut^32^, was trained to track features of the probe housing and distinct body parts of the mice (Y-maze: microdrive probe base (head), left and right ears, base of the tail, transparent linear track: forepaws, underhead/chin area). For position analysis and trial segmentation on the Y-maze, the probe base position (head position) was used as the actual position of the mouse.

### Y-maze task and trial segmentation

Mice navigated an elevated Y-maze with equally spaced arms (arm length: 45 cm, arm width: 5 cm) in search of liquid food reward (sweetened evaporated soymilk), which was dispensed from two out of three ports at the ends of each arm. The task consisted of six to eight 50-80 trial blocks, with the locations of the active ports switching in an uncued manner between blocks. The active ports were chosen randomly within each block, with no block containing the same two active ports as the previous block. Animals could not visit the same port on two consecutive trials to receive a reward, so mice learned to alternate between the active ports. For trial segmentation, we first used the *track_linearization* package^33^ to map the animal’s position on the Y-maze to a simplified one-dimensional representation with a corresponding maze arm identifier. We then defined a track “position threshold” at 70% of the length of each maze arm (31.5 cm measured outward from the center of the maze or 13.5 cm measured inward from the end of the arm). Trials were defined as periods from when the animal, after leaving a reward port, crossed the position threshold of one arm, made a turn, and crossed the position threshold of another arm without entering any other arms. The time at which an animal reached the bifurcation was defined as the time at which the animal’s linearized head position transitioned from one arm to another at the center point of the maze. Animals took ∼1.5 seconds to complete each trial (median: 1.57 s, IQR: 1.27-1.98 s, n = 24207 trials across 12 mice).

### Head angle relative to the track and time of behavioral divergence

Head angle relative to the track was defined as the angle between the animal’s head direction vector (given by the vector perpendicular to the line connecting the ears) and the midline of the track segment from which the animal approached the bifurcation. The time of behavioral divergence was defined as the time at which the animal’s head angle relative to the track significantly diverged between left and right turns. To determine the average time of behavioral divergence across sessions and animals (n = 55 sessions, 12 animals, Figure 2A, bottom panel), we calculated a Behavioral Divergence Index (BDI) as the difference between session-averaged head angles relative to the track for left and right turn trials. BDI was calculated within 200 ms windows advanced in 20 ms steps from 0.85 s before to 0.85 s after the time at which the animal reached the bifurcation. BDI significance was assessed using a permutation test: for each window, a null distribution was generated by randomly permuting left and right turn labels across sessions (10,000 permutations) and recalculating BDI from the resulting averages. Windows were considered significant only when the observed BDI was more extreme than all shuffled values (p < 1/10,000). The time of behavioral divergence was defined as the earliest time window belonging to a continuous sequence of such significant windows.

### Head-body oscillation and phase estimation

Head-body angle was defined as the angle between the animal’s head direction vector (given by the vector perpendicular to the line connecting the ears) and its body vector (given by the line connecting the midpoint between the ears and the base of the tail). To determine the average frequency of the locomotor head-body oscillation, head-body angle traces for each trial were extracted, smoothed with a Savitzky–Golay filter (50 ms, first-order) and bandpass filtered between 3 and 12 Hz (Butterworth, third-order). For all other analyses, head-body angle traces were smoothed and bandpass filtered between 4 and 8 Hz (Butterworth, third-order). Furthermore, trials were included only if they exhibited sufficient power in the head-body angle oscillation signal. To quantify this, for each trial in a given session, power spectral density was estimated using Welch’s method and the peak spectral power within the 4-8 Hz range was identified. To ensure that trials had sufficient power for spike-phase estimation, trials whose peak power fell more than one standard deviation below the mean peak power across trials in that session were excluded from analysis.

To estimate the phase of the head-body oscillation at the time of discrete behavioral or neural events (i.e., turns or spikes), we used a Hilbert transform-based interpolation approach. Head-body angle traces were first smoothed and filtered as described above. Peaks and troughs of the filtered signal were then identified (scipy.signal.find_peaks, prominence = 1° amplitude) and the instantaneous phase was obtained using the Hilbert transform. The oscillatory signal was then segmented into half-cycles (peak-to-trough and trough-to-peak). Event times were assigned a phase by identifying the half-cycle in which the event occurred and linearly interpolating the Hilbert phase within that interval. Events occurring exactly at peaks or troughs were assigned phases of 0 and π, respectively. Full oscillatory cycles, defined by successive peaks, with durations outside the expected range (i.e., cycles faster than 8 Hz or slower than 4 Hz) were excluded from analysis.

### Transparent linear track task and stepping analysis

For stepping analysis, implanted mice (n = 2) shuttled back and forth on a transparent linear track (width: 12.7 cm (x), length: 91.4 cm (y)) for liquid food reward (sweetened evaporated soymilk). Video was recorded from below using a wide-angle lens and DeepLabCut was used to track the animals’ forepaws and underhead/chin area (head position) for analyzing locomotor dynamics. Due to video distortions at the edges of the track associated with the wide-angle lens, only periods when the mouse was directly above the camera (within a 12.7 cm (x) by 32 cm (y) area) and locomoting uninterrupted were analyzed.

Forepaw swings were defined as periods when either forepaw advanced in the y-direction at speeds greater than 37.65 cm/s (3 pixels/frame). These times were then used as temporal references to examine lateral movements of the head during forepaw swings. Head position in the x-direction was filtered between 4 and 10 Hz (Butterworth, third-order). During locomotion, the head shifted laterally during each forepaw swing, such that a right forepaw swing was linked to a leftward head movement and vice-versa (Figure 1A). Notably, this same pattern (i.e., contralateral coupling between forepaw swings and head movements) has been well characterized across a variety of walking tetrapods including rats, lizards, and humans^14–16^. To quantify the direction of head movement during forepaw swings, we aligned the filtered head position in the x-direction to the middle of each forepaw swing and calculated the instantaneous change in head position (d(head x pos. filt.)/dt, Figure S1B). In both mice, left forepaw swings were associated with positive (i.e., rightward) changes in head position while right forepaw swings were associated with negative (i.e., leftward) changes in head position (Figure S1B and S1C). Furthermore, when both the forepaw and head movement signals were filtered between 4 and 10 Hz, their instantaneous frequencies were correlated across mice (Figure S1D). Thus, the left-right oscillation of the head characteristic of locomotion provides a reliable proxy for the stepping cycle in mice.

### Electrode insertion, electrode coordinates, histology and recording site assessment

In six mice, the right SC was targeted; in four mice, the left SC was targeted; and in two mice both the right and the left SC were targeted simultaneously. Lambda was used as the skull reference for all SC craniotomies. In five mice, the SC was targeted by drilling a craniotomy at anteroposterior (AP, from lambda): 0 mm, mediolateral (ML): ± 2.0 mm. The silicon probes(s) were then inserted at a 30° angle from vertical (i.e., a 30° medial tilt along the coronal plane). In six mice, the SC was targeted by a craniotomy at AP: 0 mm, ML: ± 2.5 mm and the silicon probe(s) were inserted at a 32° angle from vertical. In one mouse, the SC was targeted by a craniotomy at AP: 0 mm, ML: -1.0 mm and the silicon probe was inserted vertically with a 10° anterior tilt. Prior to implantation, the backs of all probe shanks were coated with DiI solution (Fisher, V22885) for probe track localization using a fluorescence microscope (Figure 2A, Figure S2A). The shanks were initially lowered to 0.7-1.2 mm below the surface of the brain (dorsoventral, DV) and advanced over consecutive days to reach the dSC. The dSC was targeted using DV coordinates. Once the dSC was reached, probes were advanced between sessions (days) to record from different neurons. For implantations in which the probe was inserted at an angle (n = 13 implantations from 11 mice (9 unilateral, 2 bilateral)), the average depth for the first dSC recording was DV: -1.20 ± 0.11 mm measured from the topmost channel of the probe while the average depth for the last dSC recording was DV: -2.83 ± 0.07 mm measured from the bottommost channel of the probe (Figure S2B, left panel). For the single vertical implantation, the recordings spanned from DV: -0.88 to -2.30 mm (Figure S2B, right panel).

For anatomical analysis, mice were perfused transcardially with phosphate-buffered saline (PBS) and then with 4% paraformaldehyde (PFA) in PBS. Brains were extracted, stored in 4% PFA overnight at 4°C and subsequently cut with a vibratome to 100 µm thick coronal sections. Slices were mounted in a Vectashield mounting medium containing DAPI (Vector Laboratories H1500) and fluorescence images were acquired with an Olympus MVX10 MacroView microscope. Probe tracks were identified using the DiI signal.

### Turn Direction Selectivity Index (TDSI) and turn cells

Turn selectivity was quantified using a Turn Direction Selectivity Index (TDSI), calculated as the difference between the average firing rates during right and left turns divided by their sum ((Right FR - Left FR) / (Right FR + Left FR)) over a 200 ms window centered on the time of the cell’s peak firing rate. Thus, right turn cells were defined by positive TDSI values and left turn cells were defined by negative TDSI values. To obtain firing rates, spike trains were converted to binary vectors and smoothed using convolution with a Gaussian kernel (50 ms window width). The resulting spike density functions were normalized by the kernel width to express firing rate in spikes per second and averaged across trials. To determine the time window for the TDSI calculation, we first defined a wide window of ± 0.5 s around the time at which the animal reached the bifurcation and, for each neuron, identified within this window the time of peak firing rate for either left or right turns (whichever was greater), using the neuron’s average firing rate across all arms. TDSI was then computed across each maze arm within a narrow window of ± 0.1 s (200 ms) centered on this peak firing time.

TDSI significance was assessed using a permutation test. For each neuron and maze arm, left and right turn trials were pooled together and randomly reassigned into two groups whose sizes matched the original sample sizes. TDSI was then recomputed within the narrow window centered on the same actual peak firing time. This procedure was repeated 1000 times to generate a shuffled distribution of TDSI values. For right turn cells, the p-value was defined as the fraction of shuffled TDSI values that were greater than or equal to the observed TDSI, or less than or equal to the observed TDSI for left turn cells. Turn cells were defined as neurons whose TDSI retained the same sign and remained significant (p < 0.05) regardless of which maze arm the animal approached the bifurcation from.

### Instantaneous TDSI and significant TDSI onset

We computed instantaneous TDSI across the entire trial period^6^ to determine the time at which turn cells became directionally selective (significant TDSI onset). TDSI was calculated within 200 ms windows advanced in 20 ms steps from 0.85 s before to 0.85 s after the time at which the animal reached the bifurcation. TDSI significance was assessed with a permutation test: for each window, left and right turn labels were randomly permuted across trials and TDSI was recomputed. This process was repeated 500 times to generate a null distribution for each window. Windows were considered significant when the observed TDSI was more extreme than all shuffled values (p < 1/500). Significant TDSI onset was defined as the earliest time point at which a turn cell exhibited two consecutive significant TDSI windows in the expected direction (e.g., for a left turn cell, two consecutive significant TDSI windows with negative TDSI values).

### Phase-locking analysis of turn cell activity to the head-body oscillation cycle

The phase of the head-body oscillation cycle at the time of each spike was estimated using a Hilbert transform-based interpolation approach (see *Head-body oscillation and phase estimation*). A cell’s preferred firing phase was defined as the circular mean of all spikes pooled across all trial periods (see *Circular statistics*). Preferred firing phases were also calculated separately for spikes occurring during left or right turns, and straight locomotor periods (Figure S4A and S4B). To obtain firing rates relative to the head-body oscillation cycle, spike counts were first divided by the total number of cycles and then by the average time per bin based on the average head-body oscillation frequency across animals (5.5 Hz). Phase locking of turn cell activity was assessed via a Rayleigh’s test for circular uniformity (see *Circular statistics*).

### Head-body cycles from turn

The unwrapped phase of the head-body oscillation signal was obtained for each trial via the Hilbert transform and referenced to the cycle containing the turn. Specifically, the nearest peak of the head-body oscillation signal relative to the turn was used as cycle zero by subtracting its phase value from the unwrapped signal. Preceding or following peaks were defined as cycles -1,

+1, etc. Spike times were then assigned a phase value by linearly interpolating the unwrapped phase at each spike time. Phase-aligned spike counts were then binned (bin width = π/3) and divided by the number of trials. To obtain firing rates, these spike counts per trial were further divided by the average time per bin based on the average head-body oscillation frequency across animals (5.5 Hz). Analyses were done separately for left and right turn trials.

### Straight locomotor periods

Straight locomotor periods were defined as periods in which the animal traversed the outer region of the maze, away from the bifurcation. Specifically, we defined a region comprising the outer 50% of the previously defined 70% arm segment length used for trial segmentation (i.e. from 13.5 to 29.25 cm inward from the end of the arm; see *Y-maze task and trial segmentation*). Preferred firing phases during straight locomotion were estimated using only spikes that occurred when the animal was in these regions of the maze, during both inward and outward traversals.

### Circular statistics

To quantify the relationship between rhythmic turn cell firing and the head-body oscillation cycle, we employed several circular statistical analyses. Spike times were first converted to phase (see *Head-body oscillation and phase estimation*). The preferred firing phase was calculated as the circular mean of all spike phases, determined by the angle of the mean resultant vector. To determine the strength of phase-locking and its statistical significance, we calculated a Rayleigh’s Z statistic for each cell, a standard method for assessing phase-locking between neuronal spiking and local field potential oscillations^34^. Rayleigh’s Z statistic uses the formula *Z = nR*^2^, where *n* is the number of spikes and *R* is the mean resultant length. The associated p-value of the Rayleigh’s test was estimated as *p = e^(-Z)*; this approximation quantifies the probability that the observed clustering of spikes during the stepping cycle occurred by chance, assuming a null hypothesis of uniform circular distribution. For *n* > 50, *p* = *e^(-Z)* is adequate^34^. To account for multiple comparisons across the population of 411 turn cells, a Bonferroni-corrected significance threshold was applied (a = 0.05/411). A cell was considered significantly phase-locked if its p-value was less than this adjusted alpha level (47% of turn cells, 195/411).

To identify the subpopulation of turn cells with the strongest rhythmic activity (Figure 3C), we employed a two-step filtering process. First, we identified all cells with a Rayleigh *Z* statistic meeting a nominal significance threshold of p < 0.05 (69% of turn cells, 282/411). To normalize the distribution of these values, we calculated the natural log of their Rayleigh *Z* scores (ln(*Z*)). We then defined a “high-strength” subpopulation consisting of cells whose ln(*Z*) exceeded the mean + SD of this distribution (mean: 3.11, SD: 1.45, mean + SD: 4.56). This secondary filter ensured that subsequent phase analyses focused on neurons with high phase-locking strength and most accurately defined preferred firing phases. Lastly, cells in this subpopulation with absolute TDSI < 0.1 were excluded, as very low TDSI values do not support a confident assignment to the left- or right-preferring population (this criterion excluded 1 cell). Accordingly, the high-strength subpopulation with absolute TDSI ≥ 0.1 accounted for 17% of turn cells that showed nominally significant phase-locking to the stepping cycle (49/282 cells, all of which also passed the Bonferroni-corrected significance threshold, Figure S3A).

To assess whether the preferred firing phases of left and right turn cells within this high-strength population were significantly different both during all trial periods and periods of straight locomotion (Figure 3C and 3D), we calculated the mean circular difference between the two groups. The observed difference was computed as the angular distance between the circular means of each group, mapped to the range [0, π] to represent the shortest distance on the circle. Statistical significance was assessed using a permutation test (n = 1,000 shuffles). For each shuffle, the turn cell identities (left or right) were randomly shuffled while maintaining the original sample sizes and a null distribution of angular differences was generated. The p-value was defined as the proportion of shuffled iterations where the angular difference was greater than or equal to the observed difference. Differences were considered significant if p < 0.05.

To assess differences between paired phase measurements (e.g., stepping phase at the time at which the animal reached the bifurcation for left vs. right turns; Figure 1D), we computed circular differences between paired observations and applied a Rayleigh test (*p = e^(-Z)*, as above) to assess whether these differences were non-uniformly distributed. Paired Rayleigh tests were considered significant if p < 0.05, indicating a consistent directional shift between conditions.

Correlations between distributions of preferred firing phases (Figure S4A, left vs. right turn trials, Figure S4B, all trial periods vs. straight locomotor periods) were quantified using a circular correlation coefficient (*rc*) that is robust to phase wrapping. This metric measures the degree to which deviations from the mean phase in one distribution co-vary with deviations from the mean phase in the other. Correlations were computed using the circcorrcoeff function in astropy.stats in Python. Significance was assessed using a permutation test (n = 10,000 shuffles). For each shuffle, one of the input variables was randomly shuffled and a null distribution of resulting *rc* values was generated. The p-value was defined as the proportion of shuffled iterations where the *rc* was greater than or equal to the observed *rc*.

To determine how the phase separation between left and right turn cells varied as a function of the strength of their modulation with the stepping cycle (Figure S5), we calculated the population phase relationship across a range of ln(Rayleigh Z) thresholds from 1 to 5.5 in steps of 0.5, including the ln(Z) value at the p < 0.05 significance threshold (1.13) and the mean + SD of ln(Z) values that were significant above the 0.05 level (4.56). At each ln(Z) threshold, we included all turn cells with absolute TDSI > 0.1 and whose ln(Z) exceeded that threshold and calculated, separately for left and right turn cells, the circular mean of their preferred firing phases and the mean resultant length. Phase separation was defined as the angular distance between the circular means, from 0 to π; average vector length was defined as the mean of the left and right values.

## Quantification and statistical analysis

All analyses and statistical tests were implemented using custom Python scripts (Python version 3.8.5). Statistical tests used and p-values are provided throughout the text and in figure legends.

## Acknowledgements

We thank P. Saraf, L. Ruan and Q.Y. Wu for technical support and the members of the Scanziani laboratory for helpful discussions of this project. Claude (Anthropic) was used to draft portions of the methods and figure legends, with the first author’s notes and code used as input. This project was supported by the Neuroscience Training Grant T32 (C.W.), by the Discovery Fellowship of the UCSF Graduate Division (C.W.) and the Howard Hughes Medical Institute (M.S. and L.M.F.).

## Author contributions

C.W., L.M.F. and M.S. designed the study. C.W. conducted all experiments and experimental data analysis. C.W., L.M.F. and M.S. wrote the manuscript.

## Supplemental Figures

**Figure S1:**
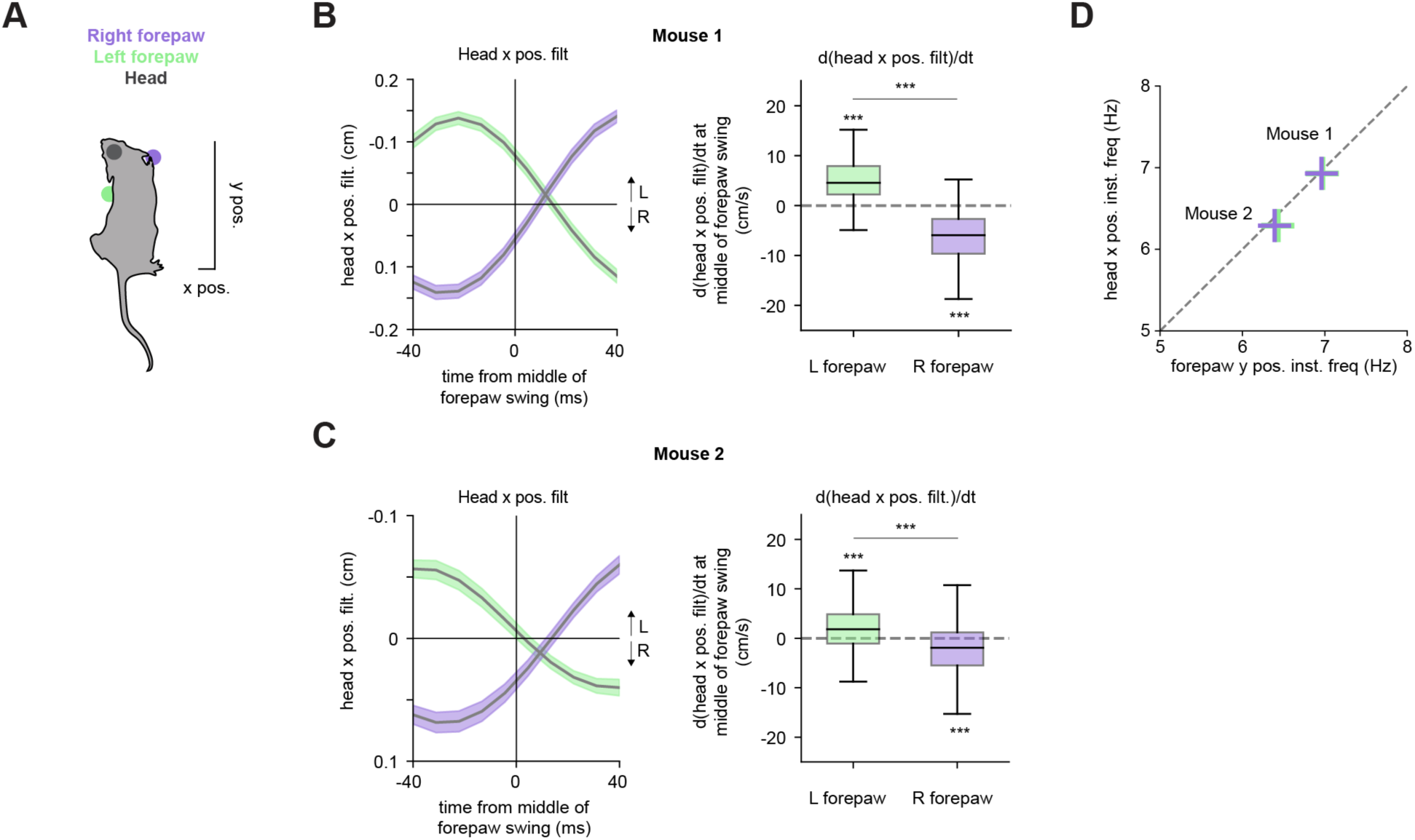
Relationship between stepping and left-right head movements. **(A)** Schematic of mouse with labels on the right forepaw, left forepaw, and head (as labeled on frames from the bottom view camera). **(B)** Left, head position along the x-axis (filtered between 4-10 Hz) for an example mouse (Mouse 1) aligned to the middle of right or left forepaw swings (color code as in (A) thick lines indicate average, shaded bands indicate s.e.m., n = 114 right forepaw swings, 105 left forepaw swings that met criteria (See Methods)). Right, rate of change of head position at the middle of each forepaw swing. Note positive (rightward) rate of change during left forepaw swings and negative (leftward) rate of change during right forepaw swings (p < 0.001 for both left and right forepaws, 1-sample t-tests compared to 0, left vs. right: p < 0.001, 2-sample t-test). **(C)** Same as in (B) but for another example animal (Mouse 2, n = 293 right forepaw swings, 326 left forepaw swings, p < 0.001 for both left and right forepaws, 1-sample t-tests compared to 0, left vs. right: p < 0.001, 2-sample t-test). **(D)** Average forepaw oscillation frequency in y position compared to average head oscillation frequency in x position. Both signals were filtered between 4 and 10 Hz.

**Figure S2:**
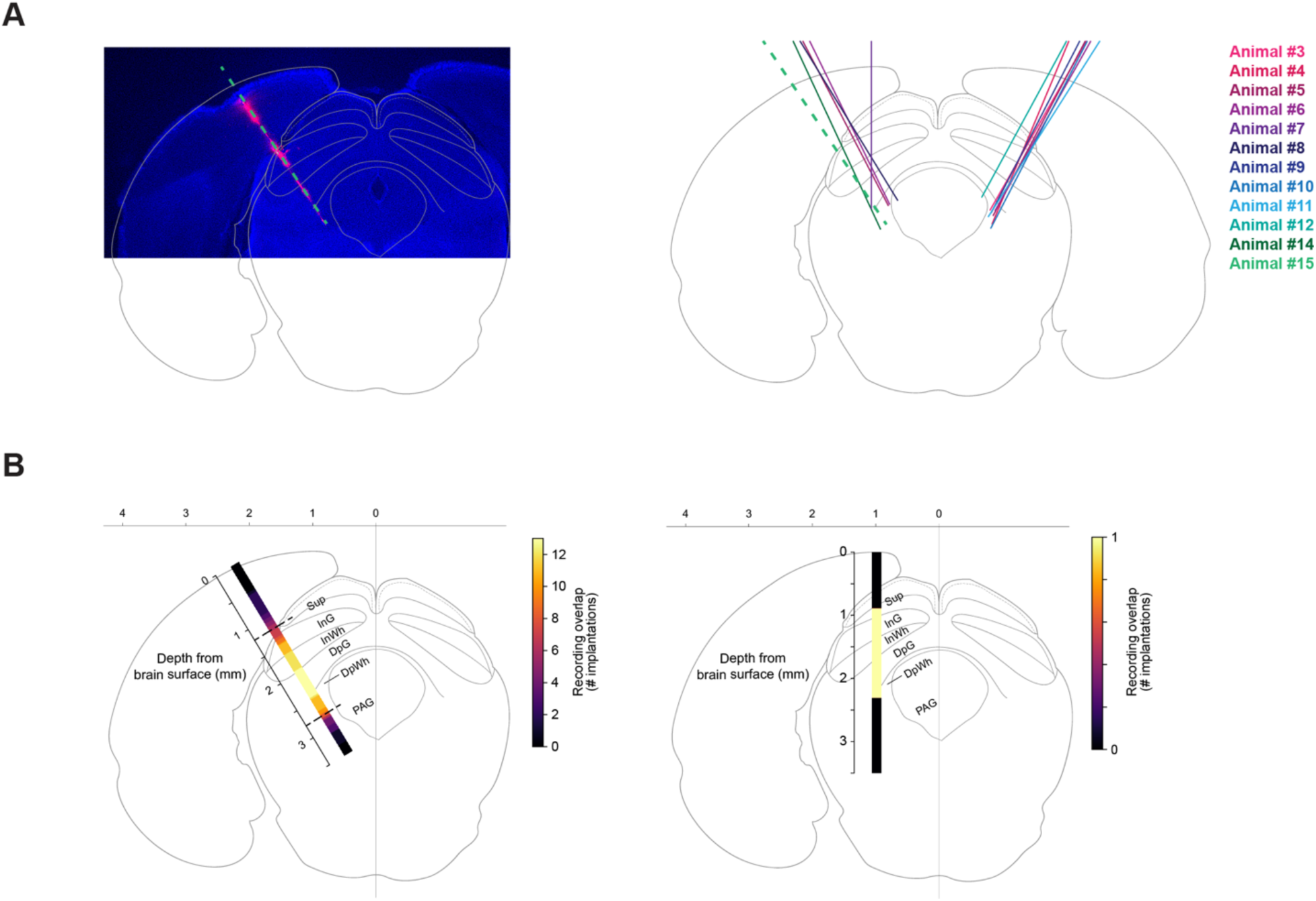
Histology and recording site analysis. **(A)** Left, example coronal section of an implantation targeting the dSC (animal #15). Probe track is marked by DiI (pink) and indicated by a dashed green line. Same as in Figure 2A. Right, probe tracks from all implantations (n = 14 implantations from 12 mice (10 unilateral, 2 bilateral)). **(B)** Left, heatmap showing the approximate span of recording depths in the SC across implantations in which the probe was inserted at an angle. Lighter colors indicate higher overlap. Dashed black lines indicate the average depths from the brain surface of the first and last dSC recordings. Note high concentration of recordings between the intermediate gray (InG) and deep white layers (DpWh). Heatmap is oriented at the average implantation angle and average ML/AP coordinates across animals (31°, ML: 2.25 mm, AP: 0 mm from lambda (-4.04 mm from bregma)). Right, heatmap showing the approximate span of recording depths for the vertical implantation. (Sup = superficial layer, InG = intermediate gray layer, InW = intermediate white layer, DpG = deep gray layer, DpWh = deep white layer, PAG = periaqueductal gray, adapted from The Mouse Brain in Stereotaxic Coordinates^35^).

**Figure S3:**
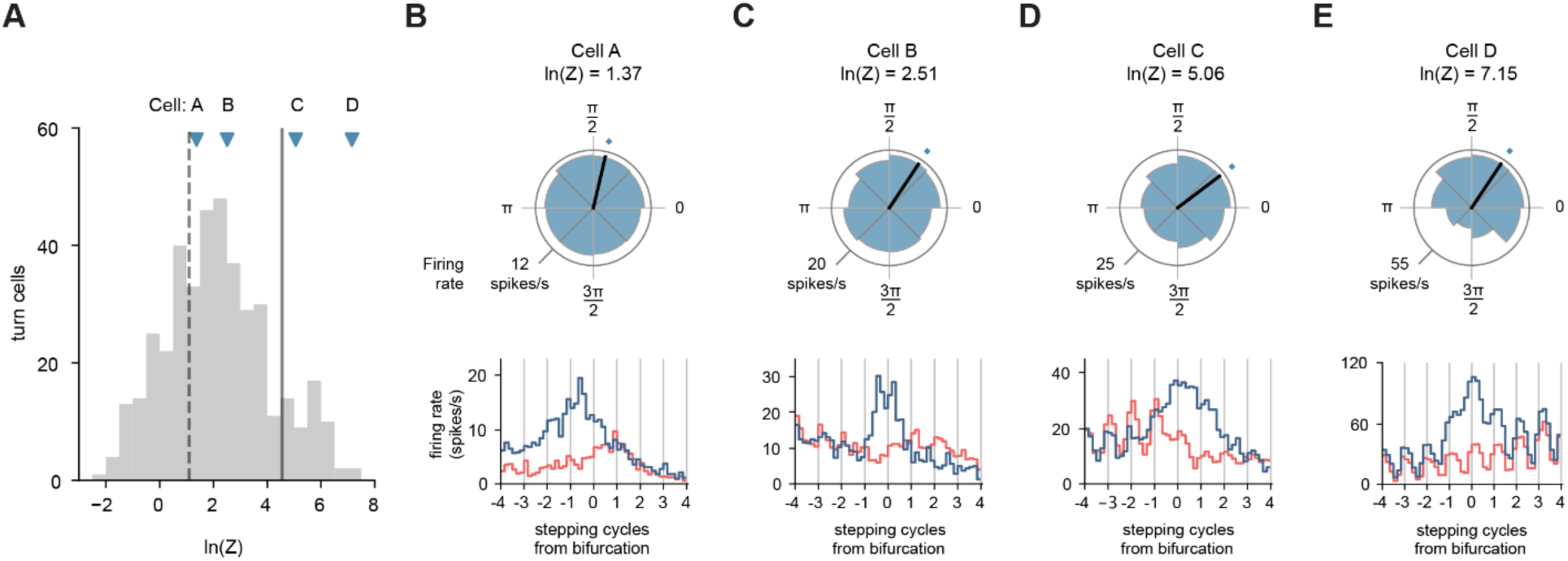
Distribution of phase-locking strength across turn cells. **(A)** Distribution of log-transformed Rayleigh’s Z values across all turn cells (n = 411 cells). Ln(Z) values to the right of the dashed vertical line are significant at p < 0.05 (n = 282 cells). Ln(Z) values to the right of the solid vertical line indicate turn cells with the strongest rhythmic activity (n = 50 cells; solid vertical line: mean + SD of ln(Z) values that are significant at the p < 0.05 level, mean + SD ln(Z) = 4.56; note that 49 cells met the absolute TDSI ≥ 0.1 criterion). Markers indicate four example rhythmically active left turn cells with increasing phase-locking strength shown in panels B-E. TDSI values for Cells A, B, C and D are: -0.53, -0.50, -0.40, and -0.55, respectively. **(B-E)** Top, polar plots of the firing of each example cell in A relative to the phase of the stepping cycle. Bottom, firing rates of each example cell aligned to stepping cycles from the bifurcation. Vertical gray lines indicate phase 0 (2π). Note the strong and reliable phase-locking of Cells C and D (ln(Z) > mean + SD) across stepping cycles compared to Cells A and B (ln(Z) < mean + SD). Cell D corresponds to Cell 3 in Figure 3.

**Figure S4:**
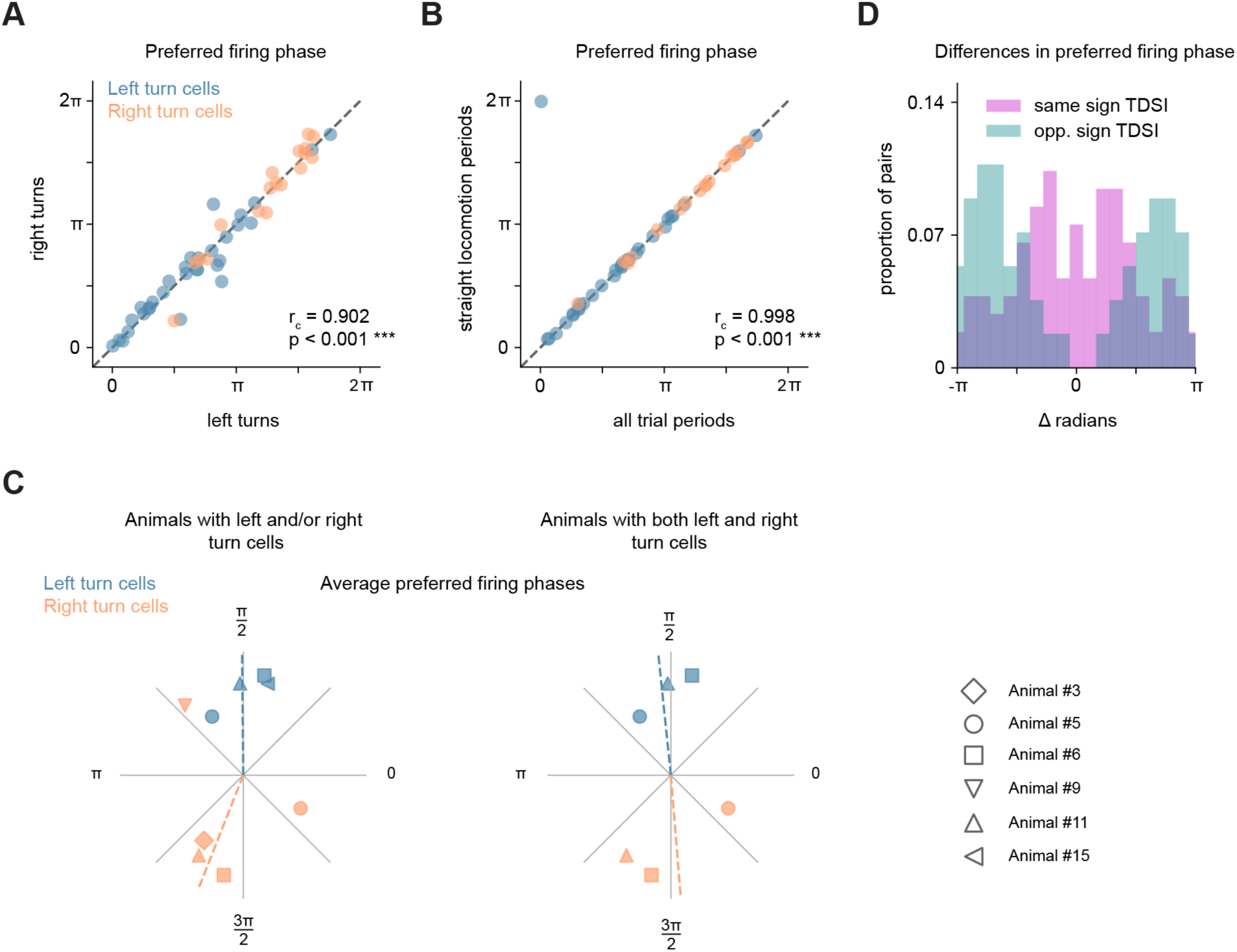
Phase relationships across turn cells whose activity is phase-locked to the stepping cycle. **(A)** Correlation between preferred firing phases of left and right turn cells during left vs. right turn trials (*rc* = 0.902, p < 0.001, permutation test). Same populations as in Figure 3C. **(B)** Correlation between preferred firing phases of left and right turn cells during all trial periods vs. straight locomotor periods (*rc* = 0.998, p < 0.001, permutation test). Same populations as Figure 3C. **(C)** Distribution of average preferred firing phases across individual animals (different symbol shapes correspond to different animals indicated to the right; same populations of cells as in Figure 3C). Left, animals contributing left and/or right turn cells (mean phase difference between left and right turn cells across animals: 2.76 rad.). Right, only animals contributing both left and right turn cells (mean phase difference between left and right turn cells across animals: 3.12 rad.). **(D)** Differences in preferred firing phases between simultaneously recorded cell pairs color coded by whether the pair had TDSI values with the same or opposite sign (same sign pairs (i.e., left turn cell vs. left turn cell), mean angle: 0 ± 0.17 radians; opposite sign pairs (i.e., left turn cell vs. right turn cell), mean angle: -3.06 ± 0.18 radians). Note clustering of same sign pairs around 0 and clustering of opposite sign pairs around ± π (same sign pairs: p < 0.01, Rayleigh test; opposite sign pairs: p < 0.001, Rayleigh test).

**Figure S5:**
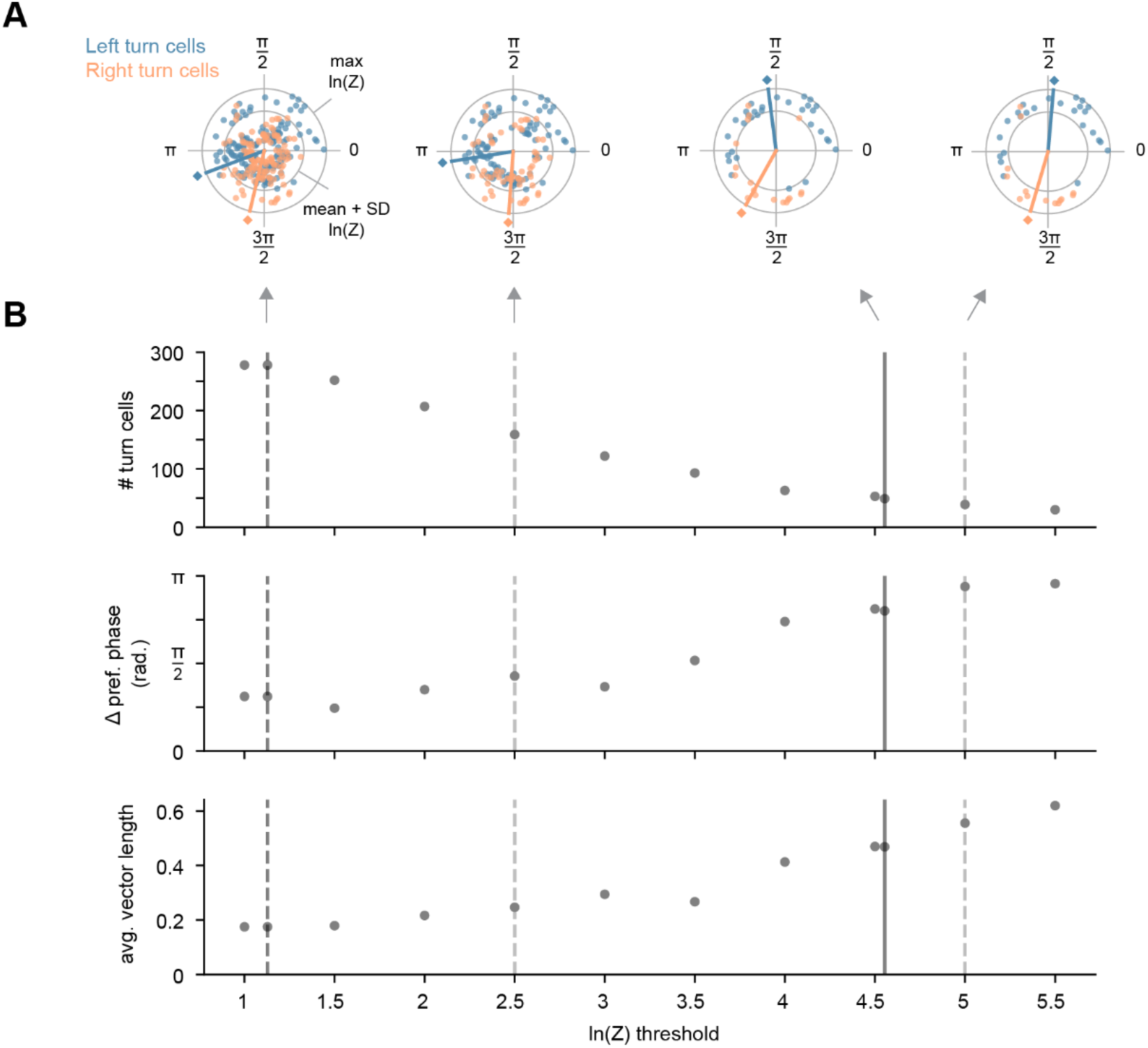
Phase separation between left and right turn cells varies with phase-locking strength. **(A)** Polar plots showing the preferred firing phases of left and right turn cells with an absolute TDSI of at least 0.1 relative to the stepping cycle. Radial axis position indicates ln(Rayleigh Z) value of each cell (outer ring: max ln(Z) = 7.20, inner ring: mean + SD ln(Z) = 4.56). Each polar plot shows the distribution of left and right turn cells at various chosen ln(Z) thresholds indicated by the vertical lines in B. **(B)** Top, scatter plot of the total number of left and right turn cells as a function of ln(Z) threshold. Dashed dark gray line indicates the ln(Z) value at the significance threshold (p < 0.05, n = 278 cells). Solid dark gray line indicates the mean + SD of ln(Z) values that are significant at the p < 0.05 level (n = 49 cells). These two dark gray lines are the same as those in Figure S3A. Dashed light gray lines indicate two additional arbitrarily chosen ln(Z) thresholds (ln(Z) 2.5: n = 159 cells, ln(Z) 5: n = 39 cells). Middle, scatter plot of the phase separation in radians between left and right turn cells as a function of ln(Z) threshold. Note that the phase separation increases toward antiphase with increasing ln(Z) thresholds. Bottom, scatter plot of the average vector length (mean resultant length, See Methods) of left and right turn cells as a function of ln(Z) threshold. Note that the average vector length increases with increasing ln(Z) thresholds.

